# Quantification of salt stress in wheat leaves by Raman spectroscopy and machine learning

**DOI:** 10.1101/2022.01.07.475441

**Authors:** Ibrahim Kecoglu, Merve Sirkeci, Mehmet Burcin Unlu, Ayse Sen, Ugur Parlatan, Feyza Guzelcimen

**Affiliations:** Department of Physics, Bogazici University, 34342, Istanbul, Turkey; Graduate School of Engineering and Science, Istanbul University, 34116, Istanbul, Turkey; Department of Biology, Istanbul University, 34134, Istanbul, Turkey; Physics Department, Istanbul University, 34134, Istanbul, Turkey

## Abstract

The salinity level of the growing medium has diverse effects on the development of plants, including both physical and biochemical changes. To determine the salt stress level of a plant endures, one can measure these structural and chemical changes. Raman spectroscopy and biochemical analysis are some of the most common techniques in the literature. Here, we present a combination of machine learning and Raman spectroscopy with which we can both find out the biochemical change that occurs while the medium salt concentration changes and predict the level of salt stress a wheat sample experiences accurately using our trained regression models. In addition, by applying different machine learning algorithms, we compare the level of success for different algorithms and determine the best method to use in this application. Production units can take actions based on the quantitative information they get from the trained machine learning models related to salt stress, which can potentially increase efficiency and avoid the loss of crops.

## Introduction

Wheat (*Triticum aestivum* L.) is one of the cereal products that can quickly adapt to environmental conditions. Besides, it is one of the most widely cultivated and produced herbal products globally thanks to the high amount of protein and carbohydrates it contains. ^1^ Wheat is a nutritious plant that meets 20-80% of the energy and protein needs of humans.^2^ Turkey produced approximately 23 million tons of wheat in 2015. This production decreased to 20 million tons in 2018. One of the significant reasons for this decrease is climate change, which affects the weather and plant growth via stressor factors such as salinity. Salinity occurs due to the accumulation of water-soluble salts in the upper part of the soil.^3^ Salinity affects the morphology, anatomy,^4^ growth, root length, and osmotic pressure of the plants.^5,6^

To determine the amount of malondialdehyde, chlorophyll, and carotenoids under optimum conditions and stress factors, biochemical methods are frequently used. The most commonly practiced technique in this type of biochemical analysis is UV-visible spectrophotometry. However, this method has some limitations in preparing, measuring, and analyzing a sample, which led scientists to seek alternatives. Recent studies utilized Raman spectroscopy to reveal the effects of salinity stress on the microstructure of wheat plants.

Raman spectroscopy is a vibrational spectral technique that gives information about the samples’ internal molecular structure illuminated with a coherent source. Since inelastically scattered photons change their frequencies as much as the molecules’ vibrational frequencies, the peaks in the spectra have intrinsic information about the sample, such as the type and amount of molecular bond vibrations.^7,8^ Raman spectroscopy has a broad use in biology due to its advantageous properties like non-invasiveness and rapidness.^9,10^ Schulz and Baranska documented the use of Raman spectroscopy in plant biology with a detailed list of tentative assignments.^11^ Studies have been carried out using Raman Spectroscopy on many plants, including fennel fruits, chamomile clusters, turmeric roots, untreated summer wheat leaves, wheat grains, and fresh soybeans. ^12–15^ Raman spectroscopy provides rapid results with little or no sample preparation. Specifically, drought stress has already been investigated employing benchtop Raman spectroscopy^13,15,16^ with the help of statistical methods.

In addition to the contributions mentioned above, the chemometric analysis helped intensely interpret the plant Raman signals.^17–19^ Cebi *et al*. detected the amino acid L-Cysteine in the wheat floor using chemometric methods such as principal component analysis (PCA) and hierarchical cluster analysis (HCA).^17^ In a food chemistry application, Raman spectroscopy and PCA were used to predict malting barley husk adhesion quality.^19^ In another application, Farber *et al*. measured the pathogen Raman spectra from Maize Kernels and distinguished healthy and diseased states with 100% accuracy using PCA and discriminant analysis (DA).^18^

We employed a machine learning-supported Raman Spectroscopy approach to determine the amount of salt stress factor the plant was exposed to. Our study employed a range of machine learning algorithms and compared their training times, prediction speeds, and success rates.

## Experimental Section

### Wheat Growth

Wheat (*Triticum aestivum* L.) seeds of Selimiye variety were obtained from Trakya Agricultural Research Institute (Edirne).

Murashige and Skoog (MS) medium was used as the basic medium for the creation of a tissue culture medium. 20 g/l sucrose (C_12_H_22_O_11_), 50 *μ*g/l 2.4-D (2.4-Dichlorophenoxyacetic Acid), and 8 g/l Agar regeneration medium were added into the medium as a carbon source, a plant growth regulator, and as a thickener, respectively.

To determine the sensitivity of wheat embryos to salt stressors, different concentrations of NaCl (50 mM 100 mM - 150 mM NaCl) were added to the regeneration medium whose content was given above. The pH values of the mixtures formed were adjusted to be between about 5.8 and 6.0. The created nutrient media were sterilized in an autoclave at 121°C at 1.2 atm pressure for 20 minutes and poured into sterile magendas in an aseptic environment in equal amounts. ^20^

For sterilization, the wheat seeds were kept in 70% ethyl alcohol for 20 seconds, passed through the sterile distilled water series 3 times, then kept in 20% commercial bleach for 20 minutes and passed through the distilled water series three times to remove the bleach. Then, the seeds were incubated in sterile distilled water for 2 hours in an oven at 35°C, and swelling was allowed. ^20,21^ The mature seed embryos were left to swell after surface sterilization. They were separated from their endosperms under aseptic conditions and inoculated into media prepared for salt stressor application. Cultures were observed at 25±2 °C temperature for four weeks in a daily illumination period of 16 hours light/8 hours dark.^20^

## Data Analysis

### Raman Spectra Pre-processing

Raman spectroscopy measurements include fluorescent profiles, as well. To analyze only the Raman peaks, one should estimate and clean this baseline profile. We determined the baseline using the method we have previously described [REF] and subtracted it from each spectra.

After correction, we applied an L2 normalization, also known as vector normalization. We calculated the vector norm of each spectrum and divided each element of the spectrum by this norm value. This operation provides a normalized spectrum whose vector norm equals one. It was shown that L2 normalization after fluorescent baseline correction is one of the most efficient methods in Raman spectroscopy. ^22^

### Machine Learning Application

After pre-processing, we trained a collection of machine learning regression models, such as linear regression, Gaussian process regression, neural network, support vector machines, and regression tree algorithms. We used MATLAB (R2021a, The MathWorks, Inc., Natick) and its regression learner app provided under the statistics and machine learning toolbox. Prior to training the models, we randomly split the data into train and test sets with a ratio of 80 to 20, respectively. While training, we used five cross-validation folds. After training, we tested the model using the aforementioned test set. The best result was given by a Gaussian process regression model with a rational quadratic kernel which is both stationary and non-degenerate covariance function (Equation 1), where r is the Euclidean distance between two values (*x, x*^*′*^) in the input domain (Equation 2), *l* is the characteristic length scale, and *α* is the scale-mixture parameter.

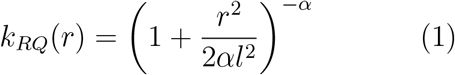

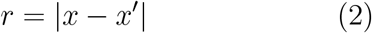

Kernel function maps the pair (*x, x*^*′*^) from input domain to IR.^23^ Basis function used was constant with a basis matrix *H* that is an n- by-1 vector of 1s, such that n is the number of observations. Initial values of the parameters were determined automatically by the regression learner app.

## Results and discussion

### Biochemical analysis using Raman spectroscopy

To determine the biochemical change per week due to different levels of salt stress, we employed Raman spectroscopy. In Figure 1 we compared the average of normalized Raman spectra corresponding to each medium salt concentrations (0, 50, 100, 150 mM) for each week. There was an expected general trend in the spectra regarding the concentration change. Although this anticipated structure was observed in the period of maturity (Figure 1c,d), there were some discrepancies in the early and late developmental periods. In the first two weeks (Figure 1a,b) plants were not yet fully grown and they showed some heterogeneity when we scanned different areas of the leaf sample. A similar but less prominent effect was also apparent in the fifth-week spectra (Figure 1e). According to the studies, when plants are exposed to longterm salt stress, they give various responses at vegetative, physiological, and biochemical levels to adapt themselves to this situation and to continue their lives under stress conditions. These responses may differ between genotypes in short-term and long-term exposure to salt stress, as in the strawberry plant.^24^

**Figure 1:**
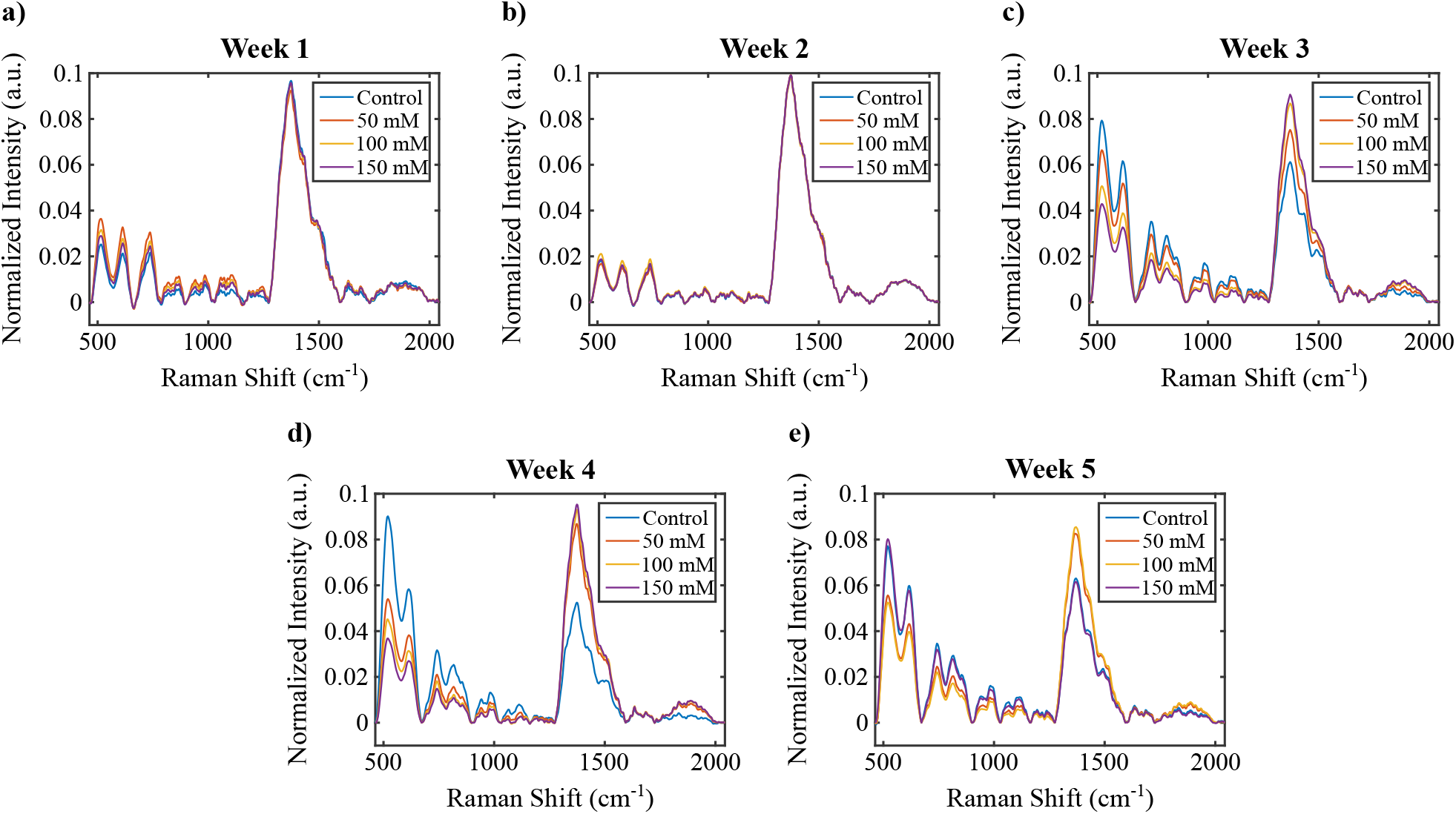
Average of normalized Raman spectra of each week.

To get a close-up to the important biochemical changes, we plotted the trend of Raman intensities regarding 522 cm^-1^, 747 cm^-1^, 855 cm^-1^, 1515 cm^-1^, and 1563 cm^-1^ bands (Figure 2). As given in Table 1, these bands correspond to cellulose, pectin, serine, carotenoid, and chlorophyll b respectively. ^11,25–28^ We observed that cellulose, pectin, and serine bands tend to decrease as the salt concentration of the medium increases, particularly for the period of maturity, which corresponds to weeks three to five. In week five, these bands have shown an increase for 150 mM concentration which contradicts the general trend. Oligosaccharides such as cellulose, pectin, and amino acids are important components that participate in cell wall formation. Therefore, in our study, cell wall components are negatively affected depending on the duration of the stress exposed to harmful metabolites such as reactive oxygen metabolites that occur due to salt stress. ^29^ The sudden increase in the fifth week suggests that the plant strives to preserve its integrity. The increase in this week suggests two possibilities. Firstly, by regulating the whole metabolism of the plant for adaptation to salt stress, it moves to a new level from the fifth week, or secondly, the measurements in the fifth week suggest that there may be an experimental error. The change in the amount of chlorophyll and carotenoid depends on the concentration of the applied salt stress and the application time, strengthening the first possibility. The decrease of these two components under normal conditions was reported in the literature.^29^

**Figure 2:**
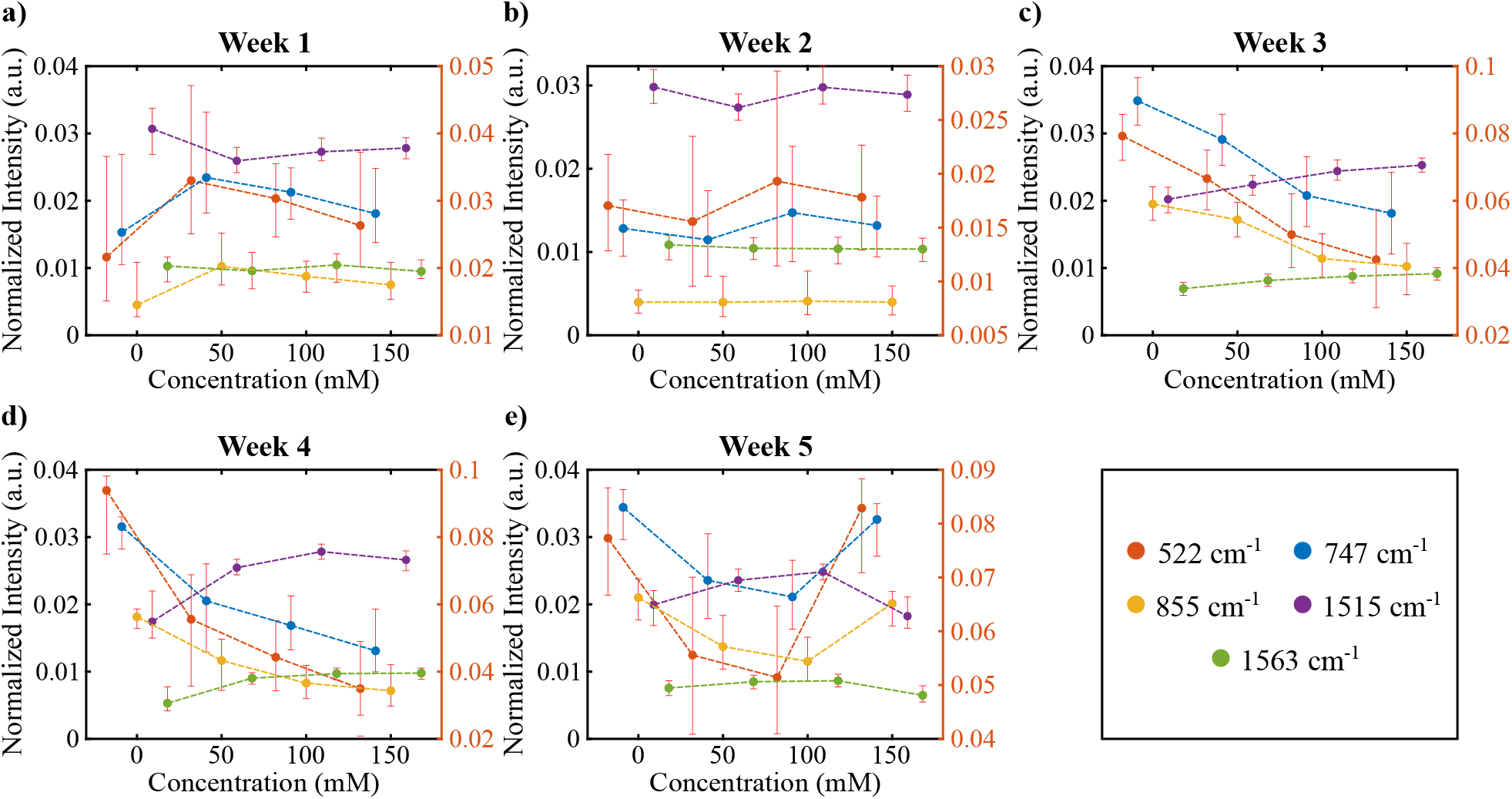
Normalized Raman intensity distributions of different concentrations for each week of plant growth. All colors correspond to different Raman shifts (cm^-1^) which is given in bottom right.

**Table 1:** Tentative assignments of Raman shift values

| Raman Shift (cm<sup>-1</sup>) | Tentative Assignments |
| --- | --- |
| 522 | $\nu(\text{C-O-C})$, cellulose<sup>25</sup> |
| 747 | $\nu(\text{C-O-H})$, pectin<sup>26</sup> |
| 855 | $\omega(\text{C}_\epsilon\text{-H})$, serine<sup>27</sup> |
| 1515 | $\nu(\text{C=C})$, carotenoid<sup>28</sup> |
| 1563 | chlorophyll b<sup>11</sup> |
| 1619 | $\delta(\text{H-N-H})$, methionine;<sup>30</sup><br>$\delta_a(\text{NH}_3^+)$, lysine;<sup>27</sup><br>$\nu(\text{C=C})$, tyrosine |
| 1646 | $\delta(\text{H}_2\text{O})$, water;<sup>26</sup><br>$\nu(\text{C=C})$, ethylene<sup>11</sup> |
| 1657 | $\nu(\text{C=O})+\omega(\text{N-H})$, amide I<sup>11</sup> |
| 1669 | $\nu(\text{C=O})+\omega(\text{N-H})$, amide I<sup>11</sup> |
| 1686 | $\nu(\text{C=O})+\omega(\text{N-H})$, amide I<sup>11</sup> |

$\nu$ - symmetric stretching, $\omega$ - wagging, $\delta$ - symmetric bending, $\delta_a$ - antisymmetric bending

We also followed the approach of Huang *et al*. to investigate further the signals related to vital markers.^31^ We modified the approach by investigating the whole Amide I region instead of taking only 1602 cm^-1^ since the samples and their responses are not the same. We utilized band component analysis, a Gaussian curve fitting procedure, to reveal the sub-components of the broad peaks measured by Raman spectroscopy. That enabled us to find the changes in the intensities and Raman shifts as the applied salt level change in particular weeks. We showed the results in Figure S2 (see supporting information). Most of the Raman intensities shown in the figure had a common decreasing trend as the weeks progressed. On the other hand, the Raman shift values have a consistent decrease only for the bands at 1619, 1669, and 1686 cm^-1^. An amide bond is a chemical bond formed between a hydroxyl group of a carboxylic group (-COOH) of one molecule and a hydrogen of an amino group (-NH2) of another molecule. Therefore, amide bonds take part in the formation of the polypeptide chain that makes up the proteins. Proteins play an important role in regulating metabolism to maintain cellular integrity under stress. ^32^ Therefore, the changes in the bands of amide bonds detected in our study show that the plant struggles with stress. The Raman bands characterize the amino acids methionine, lysine, and tyrosine, all of which participate in the structure of proteins and can be found freely in the cell. Because these amino acids participate in the regulation of the intracellular defense system depending on the stress factors such as salt stress as well as many metabolic processes in the cell.^32^ In our study, although a decrease was detected in the amount of these amino acids depending on the duration of the applied salt stress, it was determined that they increased when we evaluated the increase in the applied salt stress concentrations within themselves every week. Farhangi-Abriz and Ghassemi-Golezani (2016) ^32^ obtained similar results with our study in soybean. AlfoseaSimon *et al*., (2020) on the other hand, showed that giving amino acids such as methionine, lysine, and tyrosine exogenously to tomatoes reduced the harmful effects of salt stress in their study. ^33^

The changes in the Raman shift are a sign of molecular structure change in the plant we measured. As the number of oxidative stress products increases with the increasing salt levels, the Amide I band positions^34^ and the level of proteins during stress change.^35^ These changes were in line with the band components analysis results.

### Quantification of the acquired Raman spectra using a wide variety of machine learning regression algorithms

To quantify the salt stress level a plant endures, we trained various machine learning algorithms. We used regression models with output in terms of medium salt concentration. The output of the models can be used to determine the level of stress a plant goes through. We used five subgroups of regression models: linear regression, Gaussian process regression (GPR), neural network (NN), support vector machine (SVM), and regression tree. We gave all these models and their respective success rates, train times, and prediction speeds in Table 2. The first two weeks have shown heterogeneity. Their spectra were capriciously changing when we scanned different areas of the same sample. Consequently, we did not perform regression analysis on the spectra of the first two weeks. As a measure to determine the success rate of the models, we examined R^2^ and root mean square error (RMSE). Even though we did not consider training time and prediction speed when we determined the best-performing model, they are significant parameters, especially in realtime applications. Gaussian process regression (RMSE= 14.192-14.634, R^2^= 0.923-0.933) and neural network models (RMSE= 14.837-15.784, R^2^= 0.913-0.923) gave the best results.

**Table 2:** Comparison of the outputs of different machine learning regression algorithms. Coloring of the mean section is done for each column separately, and the performance increases as the color shifts from red to green. Abbreviations, Train: Training time (sec), Pred: Prediction speed (predictions/sec), Lin: Linear, Reg: Regression, Rob: Robust, Quad: Quadratic, Gauss: Gaussian.

|  | Week 3 |  |  |  | Week 4 |  |  |  | Week 5 |  |  |  | Mean |  |  |  |
| --- | --- | --- | --- | --- | --- | --- | --- | --- | --- | --- | --- | --- | --- | --- | --- | --- |
|  | RMSE | R<sup>2</sup> | Train | Pred | RMSE | R<sup>2</sup> | Train | Pred | RMSE | R<sup>2</sup> | Train | Pred | RMSE | R<sup>2</sup> | Train | Pred |
| Lin. Reg. | 22.972 | 0.81 | 82.516 | 1800 | 23.958 | 0.81 | 116.44 | 3100 | 23.931 | 0.81 | 83.629 | 2600 | 23.620 | 0.810 | 94.2 | 2500 |
| Rob. Lin. Reg. | 23.127 | 0.81 | 1649 | 2400 | 24.184 | 0.81 | 2713.1 | 3100 | 24.21 | 0.81 | 1922.4 | 2900 | 23.840 | 0.810 | 2094.8 | 2800 |
| GPR |  |  |  |  |  |  |  |  |  |  |  |  |  |  |  |  |
| Exponential | 14.615 | 0.92 | 2152.2 | 940 | 16.139 | 0.91 | 2371.9 | 670 | 13.149 | 0.94 | 1633 | 850 | 14.634 | 0.923 | 2052.4 | 820 |
| Rational Quad. | 14.326 | 0.93 | 2418.4 | 780 | 15.378 | 0.92 | 3642.9 | 530 | 12.873 | 0.95 | 2554.1 | 710 | 14.192 | 0.933 | 2871.8 | 673 |
| NN |  |  |  |  |  |  |  |  |  |  |  |  |  |  |  |  |
| Narrow | 17.337 | 0.89 | 2090.9 | 4400 | 16.114 | 0.91 | 3637.7 | 5400 | 13.902 | 0.94 | 2329.8 | 5300 | 15.784 | 0.913 | 2686.1 | 5033 |
| Medium | 18.632 | 0.88 | 2556.5 | 4300 | 14.411 | 0.93 | 3860.5 | 5300 | 13.013 | 0.95 | 2682.8 | 4800 | 15.352 | 0.920 | 3033.3 | 4800 |
| Wide | 17.509 | 0.89 | 3538.7 | 6400 | 14.761 | 0.93 | 5167.8 | 6400 | 12.241 | 0.95 | 3744.5 | 5700 | 14.837 | 0.923 | 4150.3 | 6167 |
| Bilayered | 16.615 | 0.9 | 2752.2 | 5200 | 15.338 | 0.92 | 4594.1 | 6900 | 14.499 | 0.93 | 3069.9 | 5200 | 15.484 | 0.917 | 3472.1 | 5767 |
| Trilayered | 16.175 | 0.91 | 2982.1 | 6000 | 14.26 | 0.93 | 4586 | 6400 | 15.321 | 0.92 | 3275.6 | 6500 | 15.252 | 0.920 | 3614.6 | 6300 |
| SVM |  |  |  |  |  |  |  |  |  |  |  |  |  |  |  |  |
| Linear | 21.3 | 0.84 | 477.31 | 140 | 24.819 | 0.8 | 856.21 | 120 | 23.328 | 0.82 | 565.5 | 130 | 23.149 | 0.820 | 633.0 | 130 |
| Cubic | 18.661 | 0.88 | 652.68 | 130 | 18.292 | 0.89 | 1175.6 | 130 | 16.745 | 0.91 | 698.65 | 160 | 17.899 | 0.893 | 842.3 | 140 |
| Quadratic | 18.704 | 0.88 | 569.08 | 130 | 18.461 | 0.89 | 1254.5 | 110 | 18.048 | 0.89 | 770.66 | 140 | 18.404 | 0.887 | 864.7 | 127 |
| Coarse Gauss | 21.382 | 0.84 | 908.3 | 170 | 26.937 | 0.76 | 1695.8 | 180 | 23.481 | 0.82 | 1068.6 | 190 | 23.933 | 0.807 | 1224.2 | 180 |
| Medium Gauss | 15.727 | 0.91 | 824.46 | 170 | 18.357 | 0.89 | 1391.5 | 150 | 14.155 | 0.94 | 895.43 | 200 | 16.080 | 0.913 | 1037.1 | 173 |
| Tree |  |  |  |  |  |  |  |  |  |  |  |  |  |  |  |  |
| Coarse | 20.472 | 0.85 | 118.48 | 5000 | 24.295 | 0.81 | 169.21 | 6900 | 22.253 | 0.84 | 131.35 | 6200 | 22.340 | 0.833 | 139.7 | 6033 |
| Medium | 21.127 | 0.84 | 71.227 | 3700 | 24.027 | 0.81 | 100.75 | 6700 | 22.477 | 0.84 | 74.088 | 5400 | 22.544 | 0.830 | 82.0 | 5267 |
| Fine | 22.94 | 0.81 | 84.19 | 3400 | 25.139 | 0.79 | 119.61 | 6000 | 22.852 | 0.83 | 86.696 | 5100 | 23.644 | 0.810 | 96.8 | 4833 |
| Bagged | 16.857 | 0.9 | 1354.8 | 4300 | 18.937 | 0.88 | 2201.1 | 5000 | 15.278 | 0.92 | 1489.4 | 5100 | 17.024 | 0.900 | 1681.8 | 4800 |
| Boosted | 18.843 | 0.87 | 1315.8 | 4500 | 21.762 | 0.84 | 2236.3 | 6100 | 19.173 | 0.88 | 1518.1 | 5400 | 19.926 | 0.863 | 1690.1 | 5333 |

In terms of mean RMSE and R^2^, the best performing model among all was rational quadratic Gaussian process regression (RQGPR). We used the test portion of the data to determine the model’s success rate. In Figure 3a-c, we showed the RQGPR model’s predictions for the test set. The median of the predictions corresponding to each true concentration value is approximately equal to the true concentration value, an indicator of a well-performing model. To make this observation more visible, we also presented residuals for each concentration in Figure 3d-f. The medians in the residual plots are all around zero. This indicates that if the age of the observed leaf is entered in terms of weeks, the model can predict the salt level of the plant.

**Figure 3:**
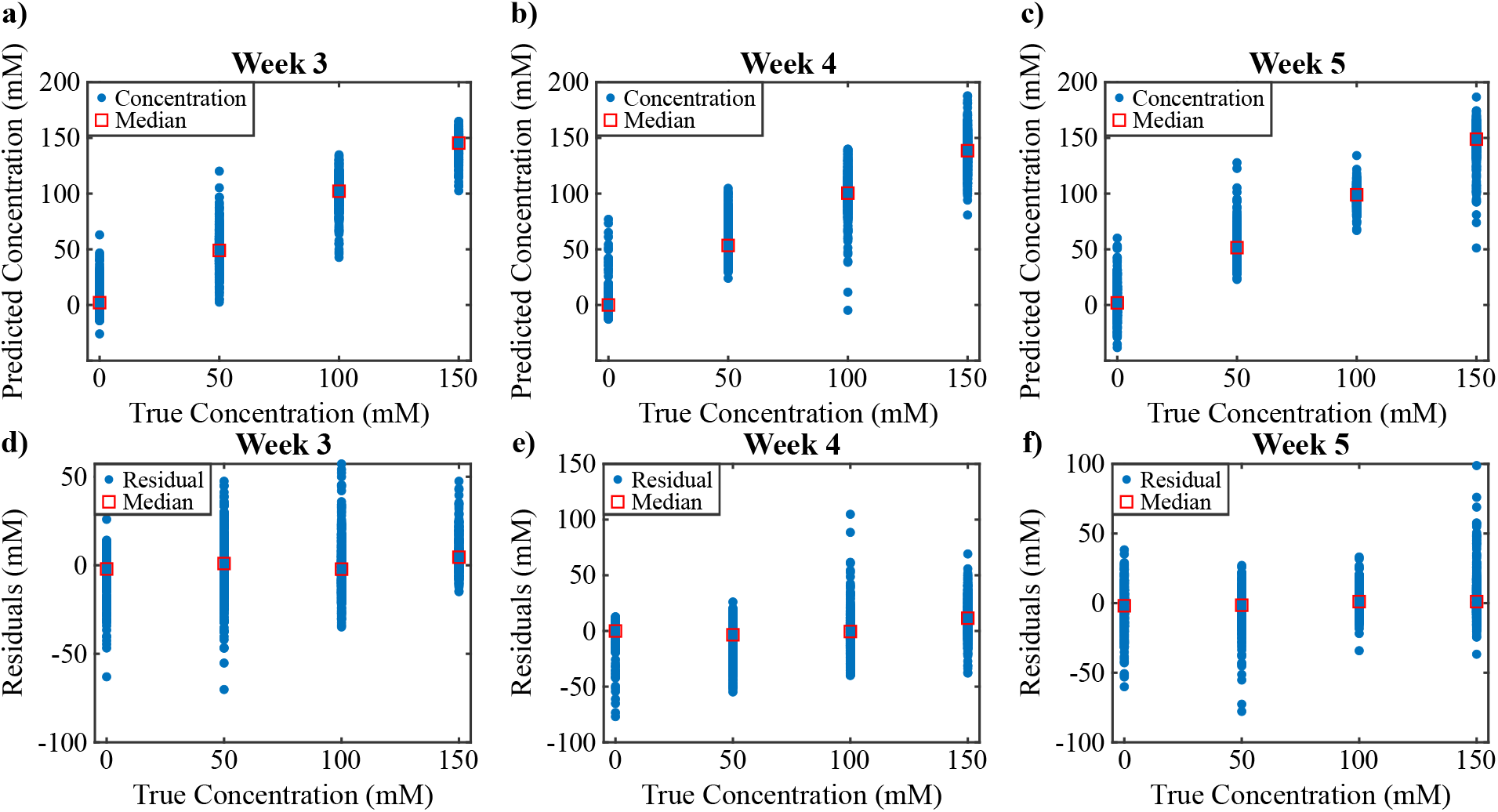
Rational Quadratic Gaussian Process Regression results (RQGPR). Out of all the models trained RQGPR has given the best results. a-c) Prediction results of the model on the test set. d-f) Residuals of the model predictions on the test set.

## Conclusion

We showed that with the help of machine learning, Raman spectroscopy could be used to quantitatively determine the level of salt stress a plant is exposed to. The shifts in the Raman spectra give precise information about how much a plant struggles from the stressor factors and how close it is to die. We also presented that models show high levels of success rates with this data; therefore, these models can be used with high accuracy. Our proposed models can quickly provide information on leaf salinity level when used in a hand-held Raman spectrometer.

## Acknowledgement

This study was supported by Research fund of Istanbul University by Project Numbers: FDP-2017-21499 and 31725. Authors thank the Ministry of Development of Turkey (Project Number: 2009K120520). We thank scidraw.io and John Chilton for the brain image in the graphical table of contents.

## Supporting Information Available

Experimental procedures and characterization data for all new compounds. The class will automatically add a sentence pointing to the information on-line:

